# Urine-HILIC: Automated sample preparation for bottom-up urinary proteome profiling in clinical proteomics

**DOI:** 10.1101/2023.07.27.550780

**Authors:** Ireshyn Govender, Rethabile Mokoena, Stoyan Stoychev, Previn Naicker

## Abstract

Urine provides a diverse source of information related to a patient’s health status and is ideal for clinical proteomics because of its ease of collection. To date, there is no standard operating procedure for reproducible and robust urine sample preparation for mass spectrometry-based clinical proteomics. To this end, a novel workflow was developed based on an on-bead protein capture, clean up, and digestion without the requirement for processing steps such as precipitation or centrifugation. The workflow was applied to an acute kidney injury (AKI) pilot study. Urine from clinical samples and a pooled sample were subjected to automated sample preparation in a KingFisher™ Flex magnetic handling station using a novel urine-HILIC (uHLC) approach based on MagReSyn® HILIC microspheres. For benchmarking, the pooled sample was also prepared using a published protocol based on an on-membrane (OM) protein capture and digestion workflow. Peptides were analysed by LCMS in data independent acquisition (DIA) mode using a Dionex Ultimate 3000 UPLC coupled to a Sciex 5600 mass spectrometer. Data was searched in Spectronaut™ 17. Both workflows showed similar peptide and protein identifications in the pooled sample. The uHLC workflow was easier to set up and complete, having less hands-on time than the OM method, with fewer manual processing steps. Lower peptide and protein CV was observed in the uHLC technical replicates. Following statistical analysis, candidate protein markers were filtered, at ≥ 2-fold change in abundance, ≥ 2 unique peptides and ≤ 1% false discovery rate, and revealed many significant, differentially abundant kidney injury-associated urinary proteins. The pilot data derived using this novel workflow provides information on the urinary proteome of patients with AKI. Further exploration in a larger cohort using this novel high-throughput method is warranted.

## 1. Introduction

The study of the human urinary proteome is becoming increasingly popular in clinical proteomics studies. Large volumes of samples are readily available with minimal invasiveness, and, in addition, soluble proteins and peptides derived from various tissues and organs are also filtered in urine, which can reflect more general health problems [1].

Plasma was long considered the best biofluid choice for biomarker discovery studies. However, the main drawback is the large protein dynamic range and therefore protein biomarkers, often expressed in minute amounts, are difficult to detect and analyse reproducibly without the use of extensive depletion and fractionation strategies [2]. In contrast, urine has a smaller dynamic range and is therefore more suitable with current analytical technologies [2]. However, urinary proteomic analysis has unique challenges, particularly in extracting soluble urinary proteins present in dilute concentrations. To date, a few groups have developed methods for urinary proteome profiling that are robust; however, reproducibility between laboratories remains a challenge. The reported methods are based on precipitation, concentration, and on-membrane protein capture [3–6], thus removing interfering compounds found in normal urine such as salts and other metabolites. The most common methods include acetone precipitation, acetone and trichloroacetic acid precipitation, ultracentrifugation, filter-aided sample preparation and various combinations thereof [3,4,7,8]. After precipitation, protein resolubilisation is often performed using urea-based buffers instead of more efficient detergent-based buffers, as detergent removal is difficult to achieve. Unfortunately, there is no consensus on the ideal sample preparation methodology for the processing of urine and this remains the individual preference of the laboratory. Furthermore, many of the current methods lack the throughput required to analyse large clinical cohorts due to bottlenecks created by steps such as precipitation, centrifugation, and buffer exchange, which are all difficult to scale and automate.

In the current study, we present a novel approach to the preparation of urinary proteome samples. The method, named urine-HILIC (uHLC), is based on direct, on-bead protein capture (from only 100 μL of urine), clean-up, and digestion. It is automated (1 to 96 samples per run) and can be easily implemented in the mass spectrometry laboratory and requires standard sample collection procedures in clinics or hospitals. The uHLC workflow was benchmarked against a urinary proteomics workflow based on on-membrane (OM) protein capture (MStern approach) [5,6], a high throughput method that can accommodate 96 samples in parallel. A 3 by 3 approach was used to evaluate both workflows, that is, 3 technical replicates processed on 3 consecutive days (n = 9 per workflow).

We then applied the novel workflow to an acute kidney injury (n = 12) pilot study to show applicability to typical proteomics research. Using the new workflow, we were able to show differentially abundant proteins and proteins known to be associated as disease markers for AKI. We show that the novel methods reported are reproducible, robust, and highly efficient and has the potential to be used routinely in future clinical urinary proteomics research.

## 2. Materials and Methods

Solvents and chemicals (MS-grade) used in the study were purchased from MERCK unless otherwise specified. All buffers were freshly prepared. Sequencing grade modified trypsin was purchased from Promega (Madison, Wisconsin, USA). MagReSyn^®^ HILIC microspheres were purchased from ReSyn Biosciences (Edenvale, Gauteng, South Africa).

### 2.1 Urine sample collection protocol and pilot study cohort

Ethics approval was received for recruitment and collection of urine samples for this study [Ethics reference: #58/2013, #271/2018 (CSIR-REC) and #120612 (WITS-HREC)]. For the development and benchmarking of the method, urine from three healthy adult men was used after informed consent (age range 26-38 years). Clinical samples were taken from unrelated patients who had been admitted to the Tshepong Hospital (North West, South Africa). All participants were HIV-positive and undergoing first-line combination ART. They were age, race and gender matched and grouped into AKI (case) and normal (control) based on their kidney function according to the guidelines set out in Kidney Disease Improving Global Outcome report [9].

Urine was collected as midstream, clean-catch into sterile urine collection containers, and transported immediately on ice to prevent degradation. No protease inhibitors were used in this study. Individual samples were centrifuged at 800 x g for 10 min to remove debris and then aliquoted and stored at -80 °C until further use.

### 2.2 Sample preparation

#### 2.2.1 Automated urine-HILIC workflow

Samples were allowed to thaw to room temperature (RT). Urine (100 μL) was mixed with 300 uL of urine sample buffer (USB: 8M Urea, 2% SDS), and sequentially reduced and alkylated using DTT (10mM v/v; 30 min, RT) and IAA (30mM v/v; 30 min, RT-dark). Thereafter, an equal volume HILIC binding buffer (30% MeCN/200mM Ammonium acetate pH 4.5) was added to the sample-USB mix (∼ 410 μL final volume). The automated KingFisher™ HILIC workflow was then followed (protocol available from), as previously described [10,11]. The automated on-bead protein capture, clean-up, and digest protocol was programmed using BindIt software v4.1 (Thermo Fisher Scientific). Briefly, magnetic hydrophilic affinity microparticles (10 μl beads/100 μl urine) were equilibrated in 200 μl of 100 mM NH_4_Ac pH 4.5, 15% MeCN. The microparticles were then transferred to the well containing the sample-USB-bind buffer mix and mixed for 30 min at RT. The captured proteins were washed twice in 200 μl of 95% MeCN and transferred to 200 μl of 50 mM ammonium bicarbonate (ABC) containing 1 μg sequencing grade modified trypsin (Promega, Madison, USA) and mixed for 2 hr at 37 °C. Finally, beads were washed in 1% TFA to elute any remaining non-specifically bound peptides. The resulting peptides (pool of digest and TFA eluate) were vacuum dried, resuspended in 2% MeCN, 0.2% FA and quantified using the Pierce™ Quantitative Colourimetric Peptide Assay (Thermo Fisher Scientific, Massachusetts, USA) according to the manufacturer’s instructions.

#### 2.2.2 On-membrane workflow based on MStern Blot

The on-membrane protein capture protocol was used to benchmark the uHLC method, since it is an emerging method in urinary proteomics for large scale clinical research [5,12]. Briefly, 100 μL of urine was mixed with 300 μL of urea sample buffer (8 M urea in 50 mM ABC). Reduction with 30 μL reduction solution (150 mM DTT, 8 M Urea, 50 mM ABC) and alkylation with 30 μL (150 mM IAA, 8 M Urea, 50 mM ABC) were carried out at RT in the dark for 30 min each. Individual wells of PVDF membrane plates (MSIPS4510, Merck Millipore) were activated and equilibrated with 150 μl of 70% ethanol/water and urea sample buffer. Samples were passed through PVDF membranes using a vacuum manifold. Adsorbed proteins were washed twice with 150 μl of 50 mM ABC. Digestion was carried out at 37 °C for 2 hr by adding 100 μl digestion buffer (5% v/v MeCN)/50 mM ABC) containing 1 μg sequencing grade modified trypsin per well. The plates were sealed with a sealing mat and placed in a humidified incubator, the resulting peptides were collected by applying vacuum and the remaining peptides were eluted twice with 75 μl of 40%/0.1%/59.9% (v/v) MeCN/FA/water. Samples were frozen at -80 °C and then dried at -4 °C using a CentriVap vacuum concentrator (Labconco, Missouri, USA). The samples were resuspended in 2% MeCN, 0.1% FA and then desalted using C18 StageTips according to the manufacturer’s instructions. Desalted peptides were frozen at -80 °C and then dried at -4 °C using a CentriVap vacuum concentrator. Finally, the peptides were then resuspended in 2% MeCN, 0.2% FA and quantified using the Pierce™ Quantitative Colorimetric Peptide Assay (Thermo Fisher Scientific, Massachusetts, USA) according to the manufacturer’s instructions.

### 2.3 LC SWATH-MS data acquisition

Individual peptide samples were analysed using a Dionex UltiMate™ 3000 UHPLC in nanoflow configuration. Samples were inline desalted on an Acclaim PepMap C18 trap column (75 μm × 2 cm; 2 min at 5 μl/min using 2% MeCN/0.2% FA). Trapped peptides were gradient eluted and separated on a nanoEase M/Z Peptide CSH C18 Column (130 Å, 1.7 μm, 75 μm X 250 mm) (Waters Corp., Milford, Massachusetts, United States) at a flowrate of 300 nl/min with a gradient of 5 – 40 %B over 30 min for benchmarking and 60 min for the pilot study (A: 0.1% FA; B: 80% MeCN/0.1% FA).

Data was acquired using data-independent acquisition (DIA) - or Sequential Window Acquisition of all Theoretical Mass Spectra (SWATH) [13], using a TripleTOF^®^5600 mass spectrometer (SCIEX, Massachusetts, USA). Eluted peptides were delivered into the mass spectrometer via a Nanospray® III ion source equipped with a 20 μm Sharp Singularity emitter (Fossil Ion Technology, Madrid, Spain). Source settings were set as: Curtain gas - 25, Gas 1 - 40, Gas 2 - 0, temperature – 0 (off) and ion spray voltage – 3 200 V.

Data was acquired using 48 MS/MS scans of overlapping sequential precursor isolation windows (variable m/z isolation width, 1 m/z overlap, high sensitivity mode), with a precursor MS scan for each cycle. The accumulation time was 50 ms for the MS1 scan (from 400 to 1100 m/z) and 20 ms for each product ion scan (200 to 1 800 m/z) for a 1.06 sec cycle time.

### 2.4 Data processing

A spectral library was built in Spectronaut™ 17 software using default settings with minor adjustments as follows: segmented regression was used to determine iRT in each run; iRTs were calculated as median for all runs; the digestion rule was set as “Trypsin” and modified peptides were allowed; fragment ions between 300 and 1 800 m/z and ions with greater than 3 amino acids were considered; peptides with a minimum 3 and maximum 6 (most intense) fragment ions were accepted. This study specific spectral library was concatenated with an in-house generated urinary proteome spectral library (using Spectronaut™ “Search Archives” feature).

Raw SWATH (.wiff) data files were analyzed using Spectronaut™ 17. The default settings that were used for targeted analysis are described in brief as follows: dynamic iRT retention time prediction was selected with correction factor for window 1; mass calibration was set to local; decoy method was set as scrambled; the FDR, based on mProphet approach [14], was set at 1% on the precursor, peptide and protein group levels; protein inference was set to “default” which is based on the ID picker algorithm [15], and global cross-run normalisation on median was selected. The final urinary proteome spectral library (peptides – 20 616, protein groups – 2 604) was used as a reference for targeted data extraction.

Default settings were used for state comparison analysis using a t-test (null hypothesis that no change in protein abundance was observed between the two groups). The t-test was performed on the log_2_ ratio of peptide intensities that corresponded to individual proteins. The p-values were corrected for multiple testing using the q-value approach to control false discovery rate [16].

### 2.5 Bioinformatic data analysis

Protein and peptide data were imported into ExPASy pI/MW [17] and GRAVY calculators. Protein data was further analysed in Spectronaut™ 17 and later exported into Microsoft Excel (v2305) to assess proteome coverage abundance scores (dynamic range assessment). Protein abundance data were analysed in ClustVis [18] and Enrichr [19] for principal component analysis and gene ontology analysis, respectively. The volcano plot was plotted by http://www.bioin-formatics.com.cn/srplot, an online platform for data analysis and visualisation. All other graphs were generated in GraphPad Prism (v9).

## 3. Results

### 3.1. Peptide yield

The workflows showed different peptide recoveries as shown in (Figure 2A). The OM workflow showed a mean peptide recovery of 0.16 μg peptide/μL urine (16.3 μg total). The uHLC workflow had a higher mean peptide recovery of 0.26 μg peptide/μL urine (26.0 μg total). For both workflows, a total of 500 μg of peptide was injected for LC-MS analysis based on colorimetric peptide assay calculations.

**Figure 1:**
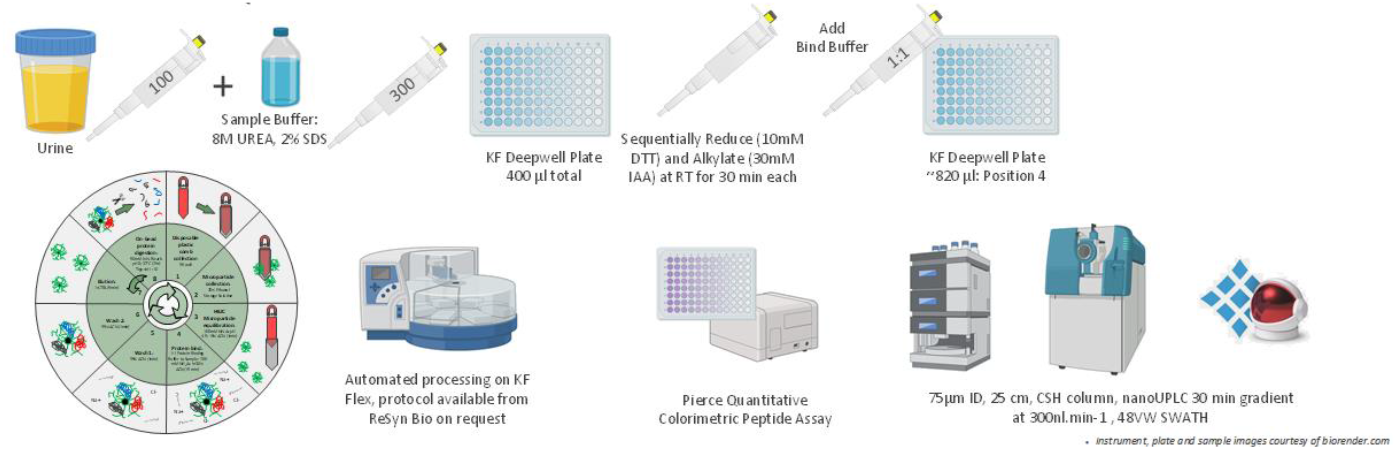
Schematic overview of the uHLC workflow.

**Figure 2:**
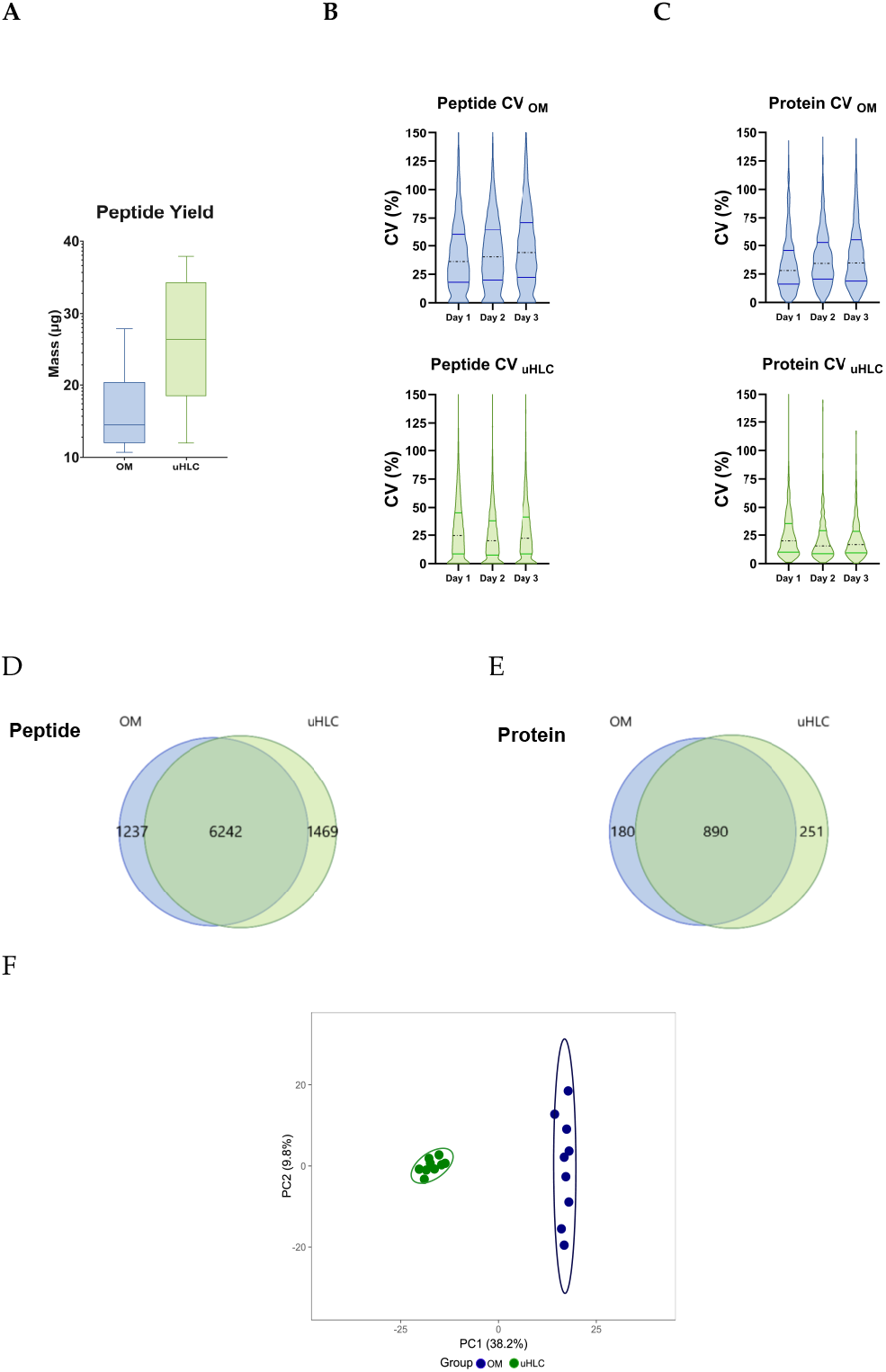
Yield and CV analysis between methods. A) Total peptide recoveries from each method. uHLC shows lower CV at the peptide (B) and protein (C) levels for all technical replicates over three days. D-E) Venn diagram showing similar protein and peptide identifications were observed. F) PCA plot shows tighter clustering of uHLC samples indicating lower CV between technical replicates.

### 3.2. Peptides and proteins identified

The uHLC workflow had higher reproducibility than the OM workflow, as shown in the lower CVs at the protein level (Figure 2B), with median CVs of 15.6% – 20% in the uHLC and 28% – 34.7% OM workflows, respectively. Similarly, at the peptide level (Figure 2C), median CVs of 20.2% – 24.7% in the uHLC and 36.2% – 44% OM workflows, were observed. PCA analysis also showed a tighter clustering of technical replicates in the uHLC workflow, indicating improved reproducibility compared to the OM workflow (Figure 2F). The workflows showed a similar total protein and peptide coverage. A large overlap was observed between the two methods, with an average of 7711 and 7477 peptides identified (Figure 2D) identified. These corresponded to an average of 1140 and 1069 protein identifications (Figure 2E).

### 3.3. Protein properties and dynamic range comparison

The protein GRAVY score, molecular weight, and isoelectric point distributions were similar between both methods, showing little to no biases (Figure 3A-C). The protein isoelectric point showed a slight difference in the number of proteins recovered below a p*I* of 8, where uHLC showed a greater overall recovery. The uHLC workflow appeared to identify more proteins (16% vs. 12%) in the lower abundance range than the OM workflow (Figure 3D).

**Figure 3:**
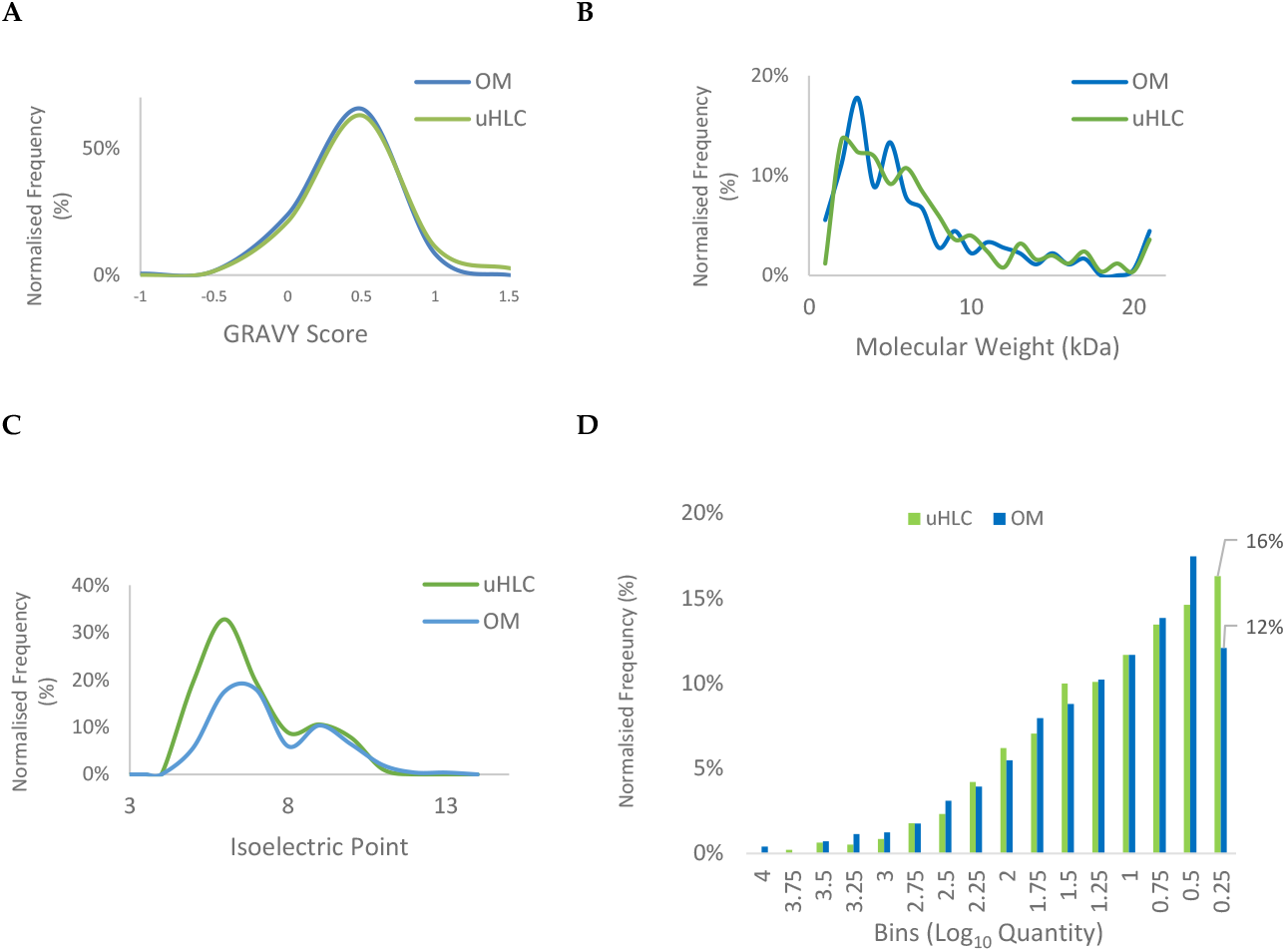
Protein properties and dynamic range comparison. A-C) Protein level analysis of GRAVY score, molecular weight distribution and isoelectric point comparing uHLC (green) and OM (blue). D) Protein abundance scores are displayed in discrete bins, from high (left) to low abundance (right).

### 3.4. Pilot study: data-independent analysis for clinical samples

The uHLC workflow was applied to a pilot cohort of 12 HIV positive patients to determine if there was a correlation between first line ARV treatment and kidney dysfunction. Participants were matched by age, race and gender and grouped into AKI (case, n = 6) and normal (control, n = 6) based on kidney function.

Data analysis of the urinary proteome revealed the presence of protein markers reported in the literature as potential biomarkers of renal dysfunction. Some known markers showed differential abundance (≥ 2-fold, q value ≤ 0.01, ≥ 2 unique peptides) between cases and controls (labelled in Figure 4A-B). The PCA analysis showed moderate clustering of the limited number of AKI and normal participants based on quantitative proteomics data (Figure 4C).

**Figure 4:**
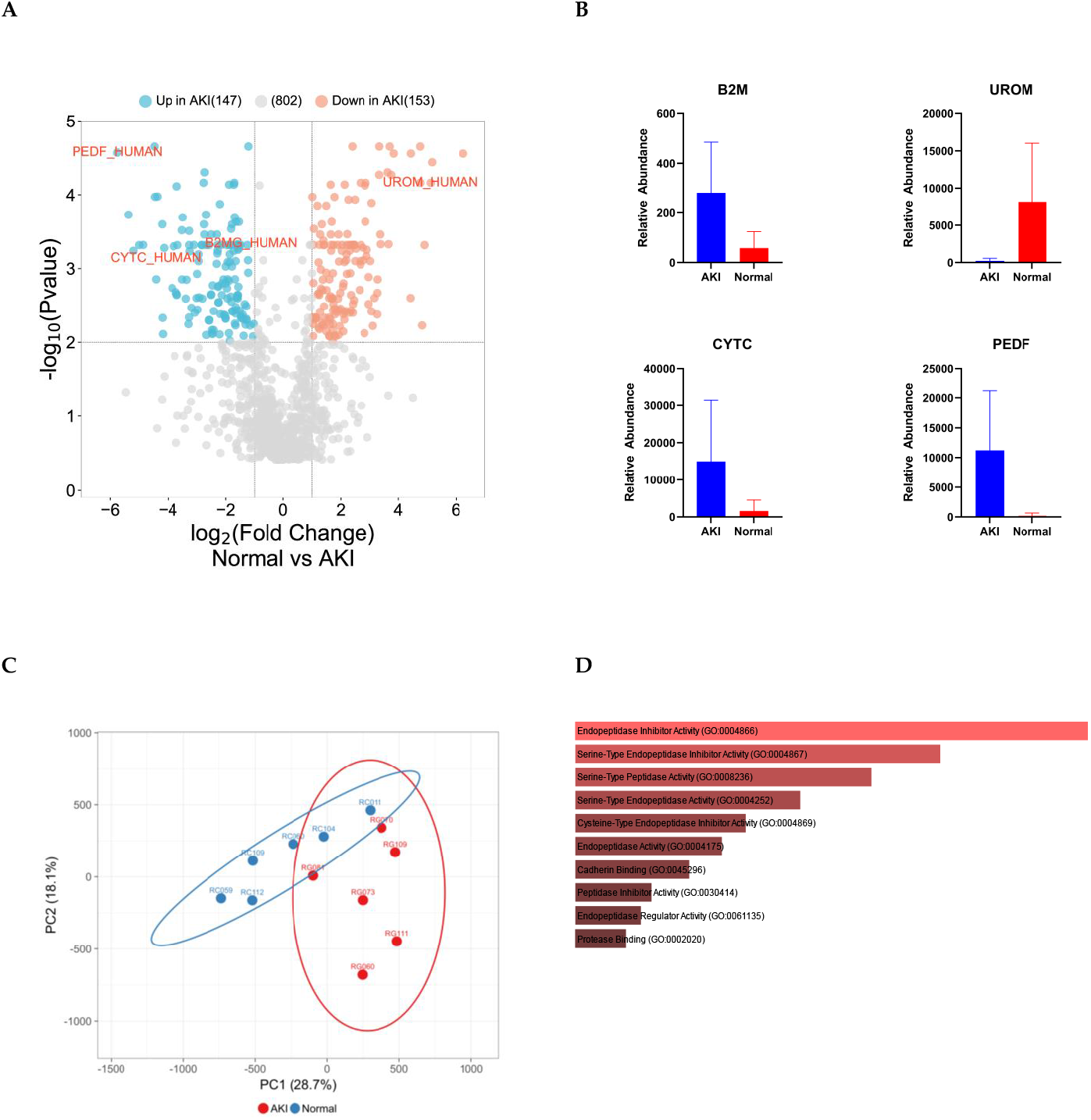
Differential analysis of pilot clinical proteomes. A) Volcano plot showing differentially abundant proteins including candidate protein markers (up in blue and down in red) and B) Known markers for kidney injury: PEDF, B2M, CYTC and UROM. C) PCA plot, the X and Y axes show PC1 and PC2 that explain 28.7% and 18.2% of the total variance, respectively. D) GO molecular function bar plot showing strong endopeptidase enrichment for the differentially abundant proteins.

## 4. Discussion

Successful biomarker studies require workflows that can be robust, easily implemented, and have high reproducibility. A generally accepted approach to urinary protein sample preparation for mass spectrometry-based proteomics is based on precipitation. After precipitation of urinary proteins, protein resolubilisation can be difficult to achieve and often requires the use of strong detergents and/or salts that are not compatible with downstream mass spectrometry analysis [20]. Urinary proteomics studies commonly use organic solvent precipitation followed by filter-aided sample preparation (FASP) as a preferred method for the isolation, clean-up, and digestion of urinary proteins [21–23], and although this is a widely used and relatively simple procedure to follow, it is a laborious process and is prone to sample loss. This is due to numerous handling steps that also have the potential to introduce sample contamination. Following FASP, samples need further processing, such as desalting and drying, before being analysed, substantially increasing cost and time and perhaps more importantly a decrease in sample recovery. These shortcomings make urinary proteome analysis using organic solvent precipitation a complicated, cumbersome, and tedious process that lacks reproducibility. Lately, there has been a drive toward large clinical cohorts, which necessitate methods that are high throughput and robust. To this end, 96-well format methods have been developed, such as MStern, which can accommodate many samples in parallel and has been shown to perform better than FASP for urinary proteomics sample preparation [6]. This is a highly successful method; however, it lacks reproducibility, mainly due to its many manual steps, and the workflow cannot be easily automated, thus limiting its use.

In contrast, we present a novel processing method, urine-HLC, that uses a small volume of urine (100 μL) mixed with urea and sodium dodecyl sulfate sample buffer with subsequent protein capture, clean-up, and on-bead digestion, using MagReSyn^®^ HILIC microspheres. The method shows performance similar to that of well-established methods such as MStern in terms of peptide and protein identification. The physicochemical properties and dynamic range of the proteins identified using both methods were similar, although some method-specific biases were observed, as expected. However, where the uHLC method was superior was in reproducibility and speed. This is largely due to the minimal handling steps and the fact that it is automated with significantly less hands-on time. Furthermore, the uHLC method appears to capture more proteins in the low abundance range, which may be important in biomarker discovery studies.

Using this methodology, we were able to confidently identify numerous markers that have been reported in the literature as potential biomarkers for various forms of kidney damage. The differentially abundant candidate markers strongly correlate with those in the literature. Beta-2-microglobulin (B2M_HUMAN) [24–26] and cystatin c (CYTC_HUMAN) [25,27,28] showed elevated urinary levels in patients with acute renal failure. A similar observation was made in kidney transplant patients who suffered rejection or postoperative renal complications in which pigment epithelium-derived factor (PEDF_HUMAN) increased in the urine after surgery [34]. Similarly, patients in our cohort who suffered kidney damage expressed higher levels of these three proteins in their urine. Uromodulin (UROM_HUMAN), the protein most abundantly expressed in the urine of healthy patients [29–31], decreased significantly in our patients with kidney injury, possibly due to tubular damage leading to decreased excretion into the tubular lumen that contains urine [32]. This finding is important in kidney injury associated with first-line ARV therapy, as it is postulated that kidney injury is due to build-up of tenofovir in proximal tubule cells leading to toxicity [33–35]. Quintana *et al*. (2009) reported similar results in which patients experiencing kidney damage expressed lower levels of uromodulin in their urine [36]. A strong enrichment of endopeptidase proteins was observed in patients with AKI, which is consistent with other studies in which these protein families showed associations with kidney injury [37].

We have developed a novel workflow, urine-HILIC, suitable for low-volume, direct, automated processing of clinical urine samples without the need for centrifugation or precipitation. The workflow shows promise for use in future urinary proteomics research and is simpler and faster than other workflows while maintaining the depth of coverage of the proteome. Furthermore, by applying the method in a pilot cohort, we were able to detect clinically relevant changes in the urinary proteome that are commonly associated with acute kidney damage. We have shown that the novel method is well suited for urinary proteome profiling and can be easily scaled for high-throughput clinical proteomics studies.

## Author Contributions

ISG and PN, designed experiments, acquired data, and wrote the first draft. RM and ISG performed wet lab experiments. ISG, SS and PN, analyzed the data. All authors edited and approved the final paper.

## Institutional Review Board Statement

The study was conducted in accordance with the Declaration of Helsinki, and approved by the WITS Human Research and CSIR Ethics Committees.

## Informed Consent Statement

Informed consent was obtained from all subjects involved in the study.

## Data Availability Statement

The mass spectrometry proteomics data have been deposited to the ProteomeXchange Consortium via the PRIDE [38] partner repository with the dataset identifier PXD043925.

## Acknowledgments

The authors would like to acknowledge the participants enrolled in the ongoing kidney injury project as well as the clinical staff at Perinatal HIV Research Unit for sample collection and patient management.

